# A Context-Aware 14-3-3 Binding Predictor Enabled by Data Augmentation, Multi-Scale Biological Features, and Protein Language Model Embeddings

**DOI:** 10.64898/2026.09.18.752707

**Authors:** Liu Liu, Zhong Wang, Xiaoqiang Huang

## Abstract

Identifying phosphorylated serine/threonine residues that mediate 14-3-3 interactions remain a major challenge in understanding phospho-dependent 14-3-3 regulation. Existing predictors largely rely on local sequence features surrounding candidate phosphosites, owing in part to the limited number of experimentally validated 14-3-3 binding sites, while broader contextual determinants of binding remain insufficiently explored. Here, we show that 14-3-3 binding sites are defined not only by local motif patterns but also by multi-scale biological context. We found that 14-3-3-binding proteins are enriched for condensation-related properties, that 14-3-3 docking sites preferentially localize to compact intrinsically disordered regions, and that protein language model embeddings provide informative representations for this task. To address both limited data availability and incomplete site representation, we developed CAMP-14-3-3 (Context-Aware Multi-scale Predictor for 14-3-3 binding), a framework that integrates multi-scale biological features with protein language model embeddings, together with a phylogeny-based data augmentation strategy and distribution-matched negative sampling. This integrated framework outperformed motif-based approaches and existing predictors on independent data. Our results support a context-aware view of 14-3-3 recognition and provide a general strategy for modeling phospho-dependent protein interactions under limited-data conditions.

**Highlights:**

* 14-3-3 binding sites exhibit distinctive multi-scale biological features, encompassing protein condensation properties, enrichment in compact intrinsically disordered regions (IDRs), and high phosphorylation propensity.
* Residue and protein-level embeddings from protein language models provide effective representations for 14-3-3 binding sites.
* Augmented 14-3-3 binding site data using an evolutionary homology-based algorithm and developed a context-aware predictor that achieves superior prediction performance.

## Introduction

The 14-3-3 protein family plays central roles in cellular regulation by interacting with phosphorylated client proteins and thereby controlling diverse processes, including cell cycle progression, apoptosis, and metabolism^1^. These interactions are classically mediated through phosphorylation-dependent binding motifs, including motif I, RSX(pS/T)XP, and motif II, RX(F/Y)X(pS)XP. However, increasing evidence indicates that many 14-3-3 binding sites deviate from these canonical consensus sequences^2^, suggesting that binding specificity is shaped not only by local motif composition but also by broader structural and biological context.

Computational identification of phosphosites that mediate 14-3-3 binding remains difficult for two major reasons. First, available training data are limited: only a few hundred experimentally validated 14-3-3 binding sites have been reported to date. Second, current representations of these sites are incomplete. Most existing predictors rely primarily on local sequence features surrounding phosphorylation sites^1–3^ and therefore are better suited to estimating peptide-level binding propensity than phosphosite-specific binding in protein context. While useful, such approaches often fail to capture the broader determinants that govern 14-3-3 binding specificity.

Recent studies have begun to point to the importance of contextual features in 14-3-3-mediated interactions. For example, 14-3-3 binding frequently occurs in intrinsically disordered regions^2^, yet such properties have not been systematically incorporated into predictive models. In addition, our previous work showed that phosphorylation within 14-3-3 binding motifs plays an important role in cellular reprogramming through regulation of biomolecular condensates^4^, suggesting that sub-organelle-level context may also contribute to 14-3-3 recognition. Together, these observations raise the possibility that 14-3-3 binding should be understood as a multi-scale, context-dependent process rather than a purely local sequence recognition problem.

Protein language models (PLMs) offer a complementary way to represent protein sequence context. Models such as ESM-2^5,6^ and ProtT5^7,8^ learn rich sequence representations from large-scale unlabeled protein data and have shown strong performance across diverse protein prediction tasks. Because PLMs can capture evolutionary and structural information without requiring extensive task-specific annotation, they are particularly attractive for problems with limited labeled data^9^. We therefore hypothesized that PLM-derived representations could provide useful complementary information for identifying 14-3-3 binding sites.

Here, we address these two challenges directly. We first improve training data through a phylogeny-based data augmentation strategy and distribution-matched negative sampling. We then develop CAMP-14-3-3 (Context-Aware Multi-scale Predictor for 14-3-3 binding) by integrating multi-scale biological features with PLM embeddings (Figure 1). Together, this framework enables more accurate identification of phospho-dependent 14-3-3 binding sites and provides a general strategy for studying protein interactions in limited-data settings.

**Figure 1.**
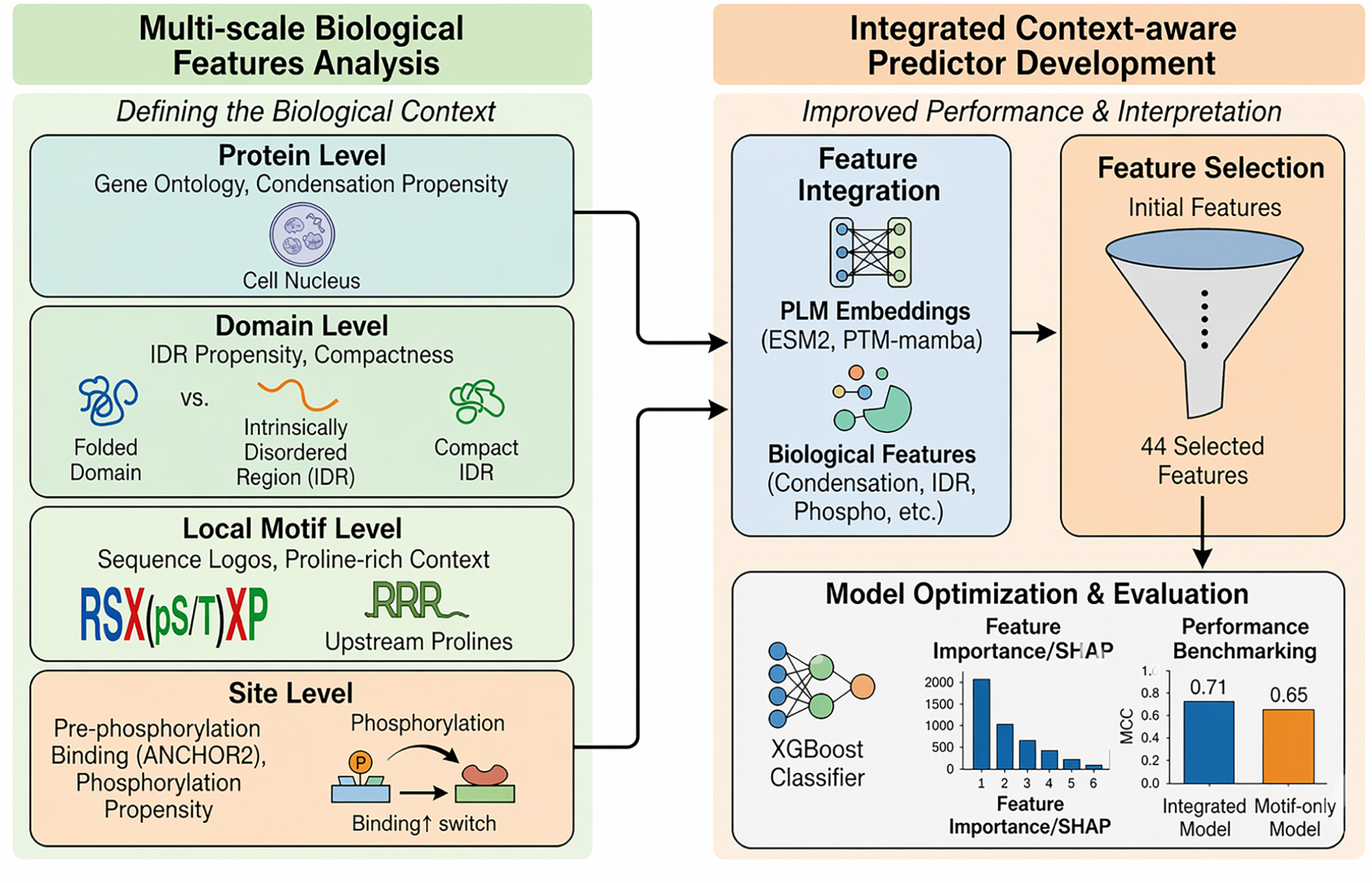
Overview of the context-aware framework for 14-3-3 binding site prediction. This figure summarizes the overall study design and prediction framework. Multi-scale biological features, including sub-organelle, protein-level, IDR/domain-level, and local motif-level properties, were integrated with protein language model embeddings to develop a context-aware predictor for phospho-dependent 14-3-3 binding sites.

## Results

### Dataset curation and augmentation for 14-3-3 binding site prediction

To systematically analyze 14-3-3 docking sites, we curated a dataset of experimentally validated 14-3-3 binding and non-binding phosphosites from published studies (Supplementary Table 1). To focus on protein-level phosphosite-dependent binding rather than peptide-level binding, we applied stringent inclusion criteria, yielding 457 14-3-3 binding positive sites from 288 proteins and 491 negative sites from 152 proteins (method) (Figure 2A). Notably, proteins containing positive and negative sites substantially overlapped, reflecting a common literature bias in which multiple candidate phosphosites are tested within the same protein (Figure 2B). Therefore, since this curated negative set may not represent the proteome-wide background distribution of serine/threonine sites, we addressed this bias through a dedicated negative sampling strategy.

**Figure 2.**
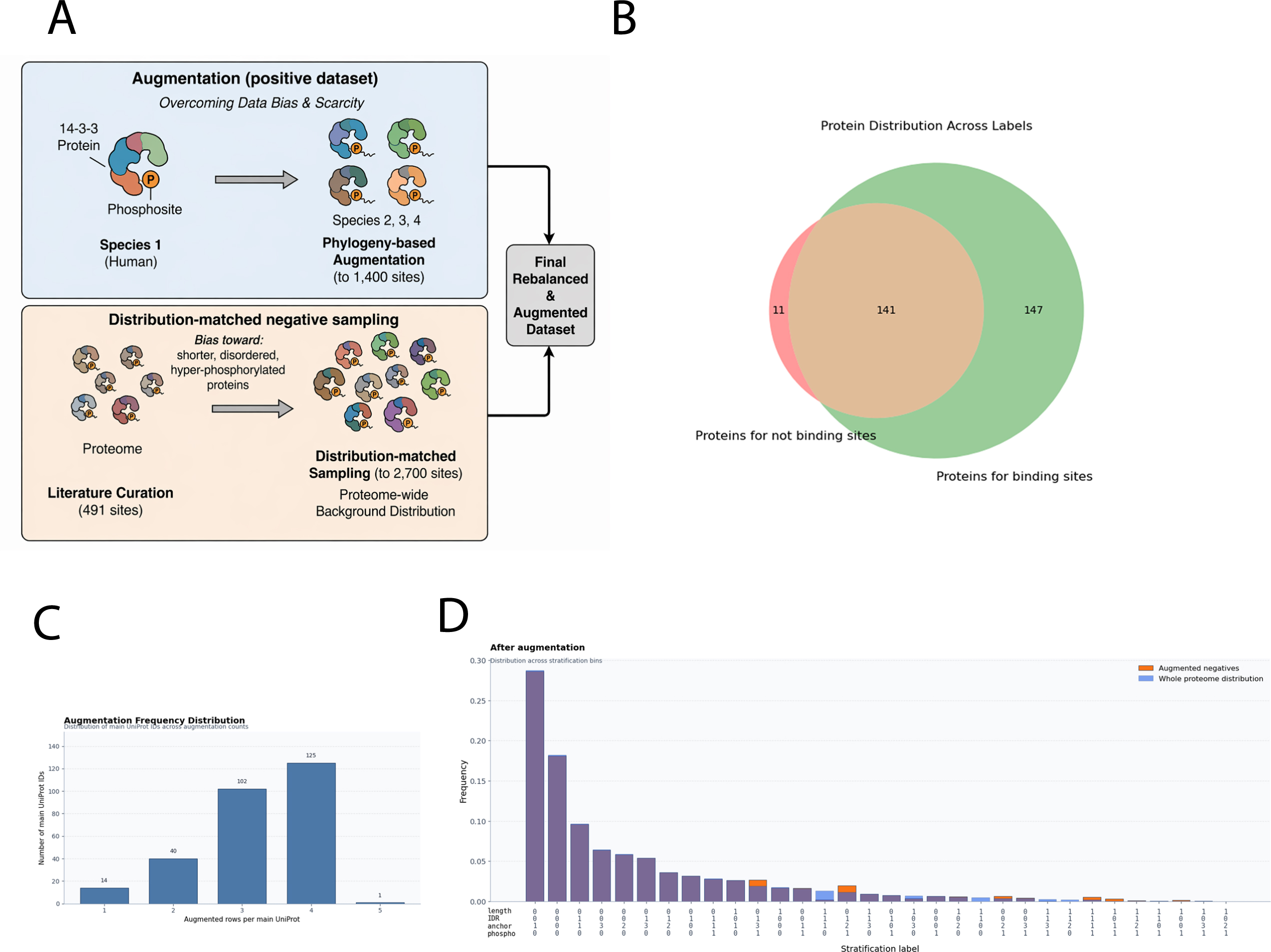
Dataset curation and augmentation for 14-3-3 binding site prediction. (A) Experimentally validated 14-3-3 binding and non-binding sites were curated and further refined through phylogeny-based positive-sample augmentation and distribution-matched negative sampling. (B) Overlap between proteins containing positive and negative 14-3-3 binding sites in the curated dataset, highlighting literature bias arising from testing multiple candidate sites within the same proteins. (C) Phylogeny-based augmentation strategy expands the positive dataset by incorporating homologous phosphosite sequences while preserving sequence diversity. (D) Distribution-matched negative sampling corrects these biases and generates a negative dataset that more closely reflects the proteome-wide background distribution of serine/threonine sites.

To overcome the limited number of experimentally validated positive sites, we developed a phylogeny-based augmentation strategy (Figure 2A). Comprehensive analysis of 14-3-3 family proteins demonstrated remarkable sequence conservation across vertebrate species, with 48 species exhibiting alignment rates > 0.6 (Supplementary Figure 1). This conservation pattern suggested functional preservation of binding interactions across species. Given the strong conservation of 14-3-3 proteins across vertebrates, we used homologous proteins carrying conserved serine/threonine residues corresponding to known human 14-3-3 docking sites as additional positive samples. This strategy expanded the positive dataset to 1,400 samples while preserving sequence diversity(method) (Figure 2C).

To generate a more representative negative dataset, we developed a distribution-matched sampling strategy based on protein-level and site-level features (Figure 2A). Relative to the human proteome, literature-curated negative sites showed substantial bias, being enriched in shorter, more disordered, and more highly phosphorylatable proteins (Supplementary Figure 1). We therefore sampled serine/threonine sites from the human proteome to match the whole-proteome background distribution, yielding a final negative dataset of 2,700 sites with no significant difference from the proteome-wide background (χ2 test, p = 1; Figure 2D).

We next tested whether the augmented and rebalanced dataset improved predictive performance. Using models trained only on local sequence features derived from the -7 to +7 phosphosite window, we found that data augmentation substantially improved performance (Table 1). On the validation set, the augmented model achieved an MCC of 0.65 compared with 0.45 for the non-augmented model. On the independent test set, the augmented model achieved an MCC of 0.65 compared with 0.59 for the non-augmented model, exceeding the previously reported value of 0.61^1^. Together, these results show that improving training data quality makes a major contribution to 14-3-3 binding site prediction.

**Table 1.** Impact of data augmentation on model performance.

|  | Model trained on non-augmented dataset | Model trained on augmented dataset |
| --- | --- | --- |
| Training dataset | 0.77 | 0.79 |
| Validation dataset | 0.45 | 0.65 |
| Test dataset | 0.59 | 0.65 |

### Multi-scale biological features of 14-3-3 binding proteins and sites

To define the biological context of 14-3-3 binding, we analyzed features across multiple scales, including sub-organelle, protein, domain, and local motif levels (Figure 1). Gene Ontology analysis revealed strong enrichment of 14-3-3-binding proteins in condensation-related functions, particularly non-membrane-bounded organelles (Figure 3A). This observation was further supported by analysis of a broader BioGrid^10^-derived set of reported 14-3-3-binding proteins. Consistently, 14-3-3-binding proteins showed elevated condensation propensity relative to the human proteome across three established predictors^11–13^ and were enriched for multiple phase-separation-related features (Figure 3B-D). These findings support a close association between 14-3-3 binding proteins and condensation-related cellular functions.

**Figure 3.**
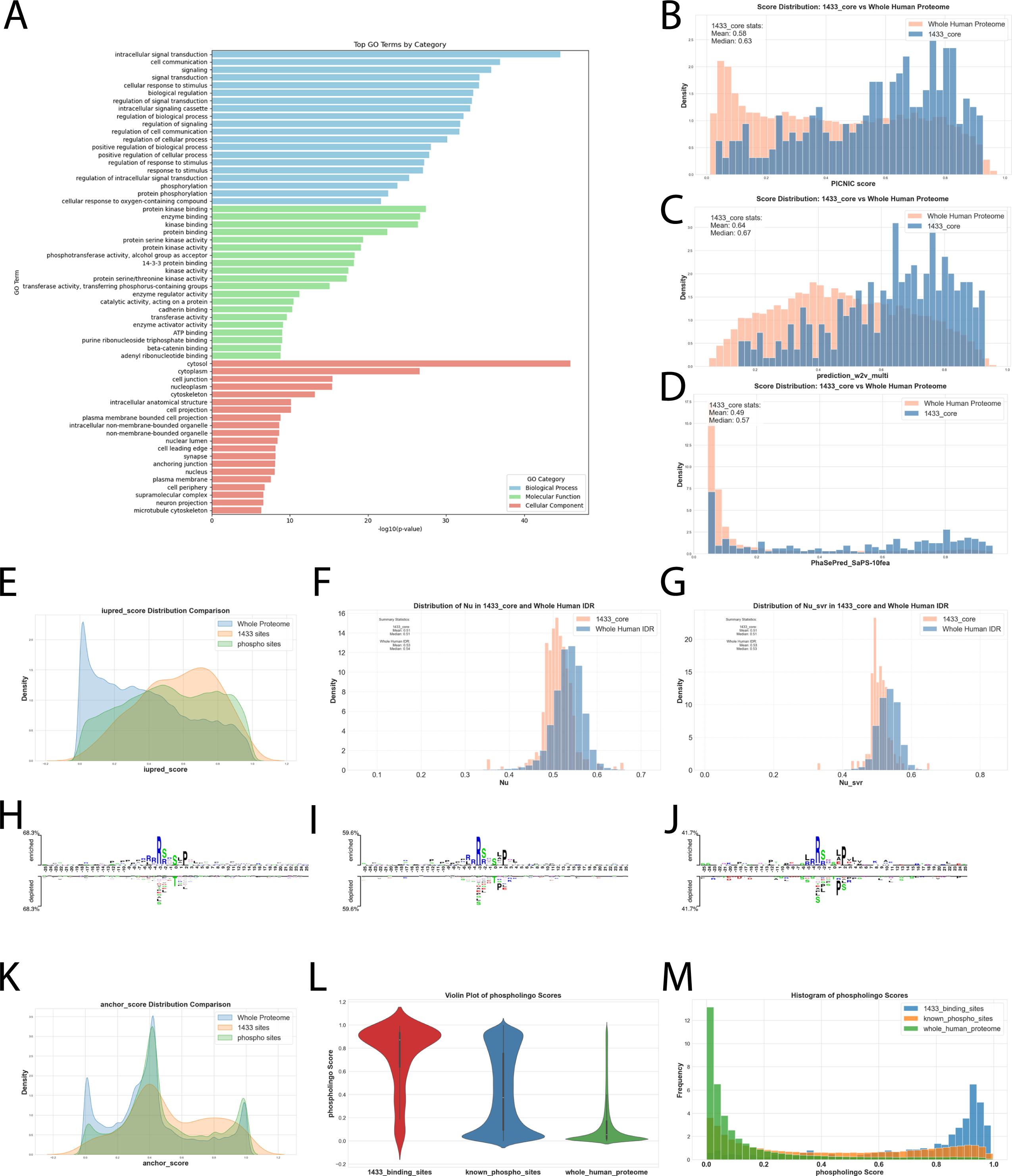
Multi-scale biological and structural features of 14-3-3 binding proteins and sites. (A) Gene Ontology analysis showing that 14-3-3-binding proteins are enriched in condensation-related functions. (B-D) Comparative analysis of the condensation propensity of 14-3-3 binding proteins using the Picnic, DeePhase, and PhaSePred predictors, demonstrating higher scores compared with the human proteome baseline. (E) IUPred2A analysis showing that 14-3-3 docking sites preferentially localize within intrinsically disordered regions (IDRs). (F–G) Analysis of IDR conformational properties showing that 14-3-3-binding sites are preferentially located in compact IDRs, as indicated by reduced Flory scaling exponent values (Nu, nuSVR). (H–J) Two Sample Logo analysis revealing characteristic local sequence patterns of 14-3-3 binding domains, including the canonical RSX(pS/T)XP motif and a proline-rich within the -12 to -7 position,indicative of condensation-prone regions. 14-3-3 positive vs 100000 sampled from whole st (Figure H), 14-3-3 positive vs all phosphorylation sites (Figure I), and 14-3-3 positive vs negative (Figure J) datasets are used for Two Sample Logo. (K) ANCHOR2 analysis showing lower pre-phosphorylation binding propensity at 14-3-3 binding sites compared with general phosphosites and proteome-wide serine/threonine sites, consistent with phosphorylation-dependent switching of interaction potential. (L–M) Phosphorylation propensity analysis via Phospholingo and show that 14-3-3 binding sites are more likely to be phosphorylated than background phosphosites and general serine/threonine sites.

Given the close link between phase separation and disorder, we next examined the structural context of 14-3-3 binding sites. Using IUPred2A^14^, we found that 14-3-3 docking sites were preferentially localized to intrinsically disordered regions (Figure 3E). Analysis of disorder-related conformational features further showed that 14-3-3-binding IDRs exhibit distinct structural properties compared with other human IDRs. In particular, 14-3-3 binding sites preferentially localized to compact IDRs, as indicated by lower Flory scaling exponent values^15^ (Nu and Nu_svr; Figure 3F and 3G), further linking 14-3-3 binding to specific disorder-related conformational states and condensation-associated behavior.

We next analyzed local sequence features of 14-3-3 binding sites. Two Sample Logo^16^ analysis identified distinct sequence patterns in 14-3-3 binding domains, including the canonical RSX(pS/T)XP motif and an enriched proline-rich upstream context (Figure 3H-J). Because 14-3-3 binding is phospho-dependent, we also examined site-level interaction and phosphorylation-related properties. ANCHOR2^17^ analysis showed that 14-3-3 binding sites had lower pre-phosphorylation binding scores than general phosphosites and proteome-wide serine/threonine sites (Figure 3K), suggesting that phosphorylation may act as a switch that enhances interaction potential. Consistent with this interpretation, 14-3-3 binding sites showed significantly higher predicted phosphorylation propensity than both general phosphosites and proteome-wide serine/threonine sites predicted by PhosphoLingo^18^ (Figure 3L-M).

Together, these results indicate that 14-3-3 binding specificity is shaped not only by local motif sequence but also by broader structural and biological context.

### Development and validation of an integrated context-aware predictor

Guided by the analyses above, we developed a Context-Aware Multi-scale 14-3-3 binding Predictor (CAMP-14-3-3) that integrates multi-scale biological features with protein language model embeddings. The biological features included condensation propensity, IDR propensity, IDR compactness, phosphorylation propensity, and local motif sequence features, whereas PLM-derived features included protein- and residue-level embeddings from ESM2 and PTM-mamba^19^.

We first compared four modeling strategies: local motif features alone, biological features combined with motif features, PLM embeddings alone, and a combined model using both PLM and biological features. Adding multi-scale biological features modestly improved performance over motif features alone, increasing validation MCC from 0.65 to 0.66 (Table 2). PLM embeddings alone were also predictive, with ESM2 outperforming PTM-mamba (validation MCC 0.61 versus 0.50). However, directly combining high-dimensional PLM features with biological features did not improve performance before feature selection. We therefore performed feature selection to identify a compact and informative representation (method). The final model retained 44 features, including ESM-derived embeddings, one-hot encoded amino acid features, and five biologically motivated features: condensation propensity, IDR propensity, IDR compactness, phosphorylation propensity, and net charge per residue (Figure 4A). These results indicate that both biological context and PLM-derived representations contribute substantially to 14-3-3 binding prediction.

**Figure 4.**
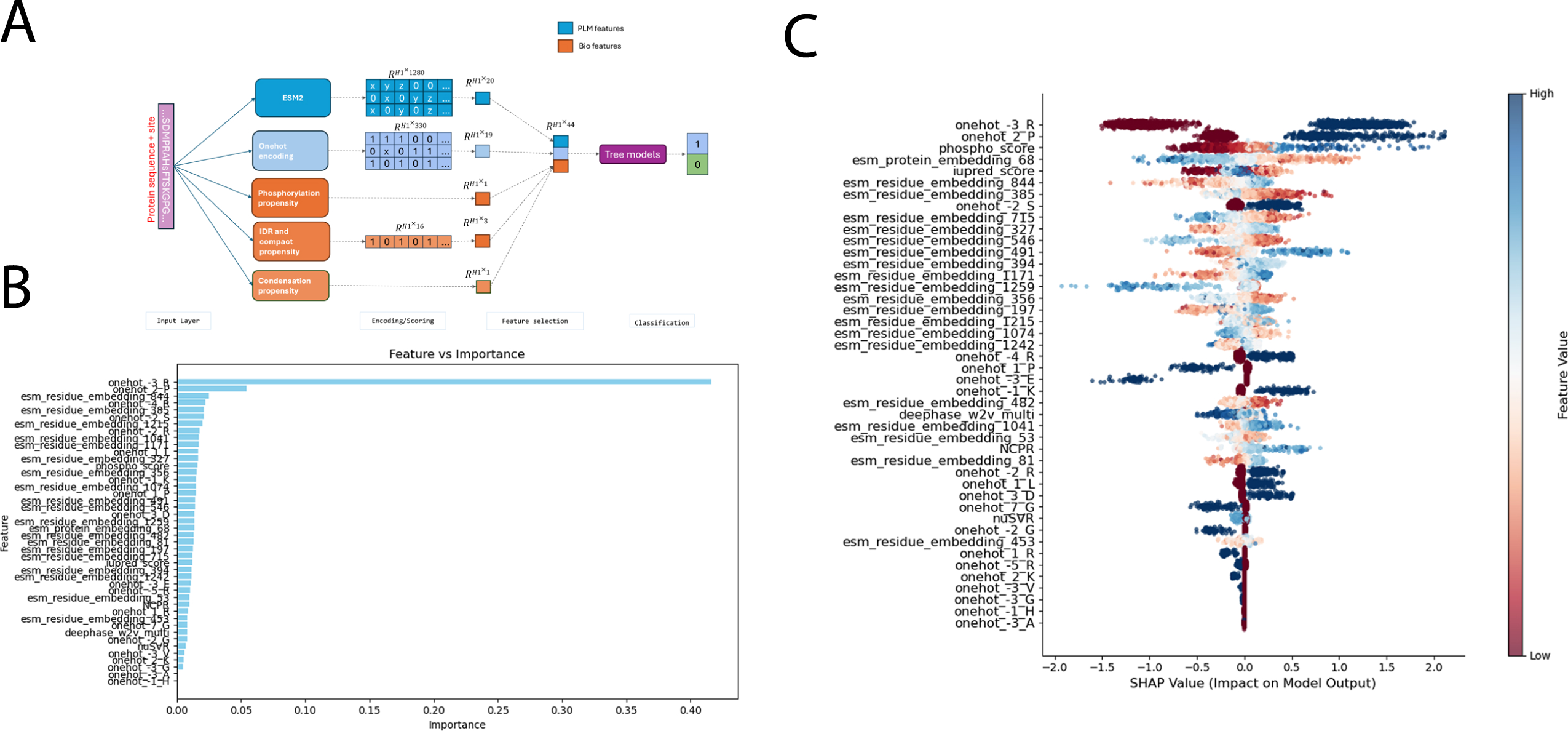
Development and interpretation of an integrated context-aware predictor for 14-3-3 binding sites. (A) Feature selection identifies a compact 44-feature representation that combines ESM-derived embeddings, one-hot encoded amino acid features, and biologically motivated features, including condensation propensity, IDR propensity, IDR compactness, phosphorylation propensity, and net charge per residue. (B) Feature importance analysis highlighting the major contributors to model performance. (C) SHAP analysis showing that both canonical motif features, such as arginine at the -3 position and proline at the +2 position, and context-related features, including phosphorylation propensity and PLM-derived embeddings, contribute substantially to prediction accuracy.

**Table 2.** Performance comparison using different features.

|  | MCC | Precision | Recall | F1-score | Accuracy |
| --- | --- | --- | --- | --- | --- |
| One-hot | 0.65 | 0.83 | 0.82 | 0.82 | 0.82 |
| Biological features + one-hot | 0.66 | 0.83 | 0.83 | 0.83 | 0.83 |
| PLM (ESM) | 0.61 | 0.82 | 0.79 | 0.79 | 0.79 |
| PLM (PTM-Mamba) | 0.50 | 0.76 | 0.75 | 0.74 | 0.75 |
| Biological features + one-hot + PLM | 0.63 | 0.82 | 0.81 | 0.80 | 0.81 |
| Biological features + one-hot + PLM<br>(selected 44 features, after fine-tuning) | 0.71 | 0.86 | 0.86 | 0.86 | 0.93 |

We next optimized the final XGBoost^20^ classifier (Table 3) and interpreted model behavior using feature importance and SHAP^21^ analyses. The most influential features included arginine at the -3 position, proline at the +2 position, and phosphorylation propensity, consistent with established determinants of 14-3-3 binding. Several ESM-derived embedding dimensions also contributed strongly, further supporting the value of PLM-based representations (Figure 4B-C).

**Table 3.** Fine-tuning of XGBoost hyperparameters through grid search.

| Hyperparameter | Range |
| --- | --- |
| n_estimators | [100, 300, 500, 1000] |
| learning_rate | [0.01, 0.05, 0.1] |
| max_depth | [2, 3, 4] |
| gamma | [0, 0.1, 0.5] |
| reg_alpha | [0, 0.1, 0.5] |
| reg_lambda | [0, 0.5, 1] |

Finally, we benchmarked our integrated predictor against existing state-of-the-art tools using an independent test dataset. For comparison, we also trained a motif-only baseline model using the same -7 to +7 local window strategy as prior methods^1^. On the independent test set, the motif-only model achieved an MCC of 0.65, whereas the integrated model achieved an MCC of 0.71 (Table 4). These results demonstrate that integrating multi-scale biological features with PLM-derived embeddings substantially improves the prediction of 14-3-3 binding phosphosites.

**Table 4.** Comparative performance evaluation on independent test set.

|  | MCC | Precision | Recall | F1-score | Accuracy |
| --- | --- | --- | --- | --- | --- |
| One-hot model | 0.65 | 0.84 | 0.81 | 0.81 | 0.81 |
| Method in this paper | 0.71 | 0.86 | 0.84 | 0.84 | 0.84 |

### Proteome-Scale Prediction of 14-3-3-Binding Proteins and Sites Across the Human Proteome

We applied CAMP-14-3-3 to the complete human proteome and systematically evaluated the 14-3-3-binding potential of every serine/threonine (S/T) site within each protein sequence. This large-scale prediction generated a comprehensive atlas of candidate 14-3-3-binding proteins and putative phosphor-dependent binding sites across the human proteome. The resulting dataset substantially expands the utility of our framework beyond single-protein analysis and provides a valuable resource for downstream functional studies and experimental prioritization of candidate 14-3-3 regulatory interactions.

### Flask-based 14-3-3 predictor implementation

We developed a lightweight Flask-based implementation of the 14-3-3 predictor and provided the source code on GitHub. The tool is designed to support local execution or user-side deployment and includes three prediction modes: single-site prediction, automatic scanning of candidate S/T residues, and sequence-level binding classification based on the highest-scoring site. The implementation reports predicted binding labels, probabilities, and relevant site information, enabling reproducible analysis of potential 14-3-3-binding phosphosites from protein sequences.

## Discussion

In this study, we systematically characterized 14-3-3 binding sites across multiple biological scales, including motif, domain, protein, and sub-organelle levels. Our results show that 14-3-3 binding specificity is related not only by local sequence patterns but also by broader structural and biological context. In particular, we found that 14-3-3-binding proteins are strongly associated with condensation-related properties and that 14-3-3 docking sites preferentially localize to compact intrinsically disordered regions. By integrating these multi-scale biological features with phosphorylation-aware representations and protein language model (PLM) embeddings, we developed a context-aware 14-3-3 binding predictor that outperformed existing approaches.

Our findings highlight the importance of incorporating biological knowledge into predictive modeling, especially for problems with limited training data. Most existing 14-3-3 binding predictors rely primarily on local sequence features surrounding candidate phosphosites, which is partly due to the scarcity of experimentally validated data. However, 14-3-3 binding is a protein-context-dependent process, and local motif information alone is insufficient to fully capture its specificity. By integrating features related to condensation propensity, intrinsic disorder, IDR compactness, and phosphorylation potential, we obtained a biologically more relevant representation of 14-3-3 binding sites and improved predictive performance. These results suggest that biologically informed feature design remains highly valuable even in the era of large pre-trained models.

We further found that PLM-derived embeddings provide complementary information for 14-3-3 binding prediction. Although PLMs are trained in a general sequence modeling setting, their latent representations capture information relevant to this downstream task. At the same time, our results indicate that not all embedding dimensions are equally informative, and that combining selected PLM features with biologically motivated features yields the best performance. This observation is particularly relevant for biological problems with limited sample sizes, where few-shot learning alone may not be sufficient and task-specific biological priors remain essential.

Another important contribution of this work is the explicit treatment of data scarcity and sampling bias. Because experimentally validated 14-3-3 binding data are limited and literature-derived datasets are biased toward proteins already known to bind 14-3-3, careful data curation is critical. By developing a distribution-matched negative sampling strategy and a phylogeny-based augmentation strategy, we improved both the representativeness and the size of the training dataset. These results emphasize that, for phosphosite-centered prediction tasks, advances in model performance depend not only on algorithm design but also on careful dataset construction.

Overall, this study provides a more comprehensive framework for understanding and predicting 14-3-3 binding sites. Rather than treating 14-3-3 recognition as a purely local sequence problem, our work supports a context-aware view in which binding specificity emerges from features spanning multiple biological scales. More broadly, our results illustrate how integrating biological knowledge, data-centric strategies, and PLM-derived representations can improve modeling of phospho-dependent protein interactions under limited-data settings.

One limitation of the current study is that the benchmark dataset remains modest in size despite augmentation, and additional experimentally validated phosphosites will further improve model generalizability. Future work may also extend this framework to isoform-specific or condition-specific 14-3-3 interactions.

## Methods

### Benchmark dataset construction

Experimentally verified 14-3-3 binding sites were collected from the literature, and detailed information on the raw dataset is provided in Supplementary Table 1. To focus specifically on protein-level phosphosite-dependent 14-3-3 binding rather than peptide-level binding, we applied stringent curation criteria. Positive samples were defined as phosphosites for which S-to-A or T-to-A mutation abolished 14-3-3 binding, as verified by immunoprecipitation. Sites supported only by peptide-level binding evidence were excluded.

Negative samples were defined as phosphosites for which single S-to-A or T-to-A mutation did not disrupt 14-3-3 binding, as well as sites for which phosphomimetic S/T-to-D/E mutation retained binding. After curation, the benchmark dataset contained 457 positive sites from 288 proteins and 491 negative sites from 152 proteins. Because proteins containing positive and negative sites substantially overlapped, suggesting bias in literature-derived negatives, additional augmentation and negative resampling procedures were applied as described below.

For model development, the final augmented dataset contained 1,400 positive samples and 2,700 negative samples. The dataset was randomly divided into training and test sets at a ratio of 4:1. To avoid information leakage, homologous augmented samples derived from the same original protein were assigned exclusively to either the training set or the test set. In addition, an independent dataset was compiled from newly collected literature cases and the experimental results reported in this study. This independent dataset had not been used in previous models and contained 264 14-3-3 sites after augmentation. Detailed information for all datasets is provided in Supplementary Table 1.

### Data augmentation

To address the limited number of experimentally validated 14-3-3 binding sites, we first developed a phylogeny-based augmentation strategy to augment positive data points. Because 14-3-3 proteins are highly conserved across vertebrate species, homologous proteins carrying conserved serine/threonine residues corresponding to known human 14-3-3 docking sites were considered candidate positive samples. Specifically, reciprocal best-hit homologs and their isoforms were identified from UniRef90 across 47 species. Candidate sequences were retained only if they contained conserved serine/threonine residues at positions corresponding to known human binding sites. To reduce redundancy, sequences were clustered using MMseq2 with a similarity threshold of 0.85, and only representative sequences were retained for subsequent analysis. Candidate homologous sites were further filtered using an existing 14-3-3 binding predictor^1^, and only high-confidence predictions were kept. To avoid overrepresentation of any single protein, no more than six homologous instances were retained per original protein. This procedure expanded the positive dataset from 457 to 1,400 samples while preserving sequence diversity. Because literature-curated negative sites are typically derived from proteins already known to bind 14-3-3, they may not represent the background distribution of non-binding serine/threonine sites across the proteome. To construct a more representative negative dataset, we developed a distribution-matched sampling strategy based on protein-level and site-level characteristics to augment negative data points. The features used for matching included protein length, intrinsically disordered region (IDR) content, anchor-site abundance, and phosphorylation propensity. Condensation-related scores were deliberately excluded from this procedure so that their discriminative value could be evaluated independently in downstream modeling.

For each feature, literature-curated negative sites and proteome-wide serine/threonine sites were compared by binning the values and estimating their joint distributions. The literature-curated negatives were enriched in shorter, more disordered, and more highly phosphorylatable proteins, serine/threonine sites were sampled from the human proteome to match the overall whole-proteome feature distribution. During this process, proteins with documented 14-3-3 interactions and sites predicted as likely binders by existing tools were excluded. Statistical validation confirmed no significant difference between the sampled negative set and the whole-proteome background distribution (χ2 test, p = 1). The final negative dataset contained 2,700 samples.

### Workflow of the 14-3-3 predictor

The overall workflow of the 14-3-3 predictor is illustrated in Figure 4. The model takes two inputs: a full-length protein sequence and a candidate serine/threonine site. After preprocessing, sequence-derived representations are generated using two feature modules: a multi-scale biological feature module and a protein language model (PLM) feature module. These features are then combined and used to train an XGBoost classifier, which outputs the predicted probability that the candidate phosphosite mediates 14-3-3 binding.

### Feature encoding schemes

Two groups of features were used to encode the input sequences: multi-scale biological features and PLM-derived features. The multi-scale biological features were selected to remain computable directly from sequence, enabling application to proteins lacking extensive experimental annotation. These included local motif sequence features based on one-hot encoding of the -7 to +7 window surrounding the candidate site, condensation-related features derived from a retrained DeePhase model, IDR propensity features from IUPred2A, IDR compactness features represented by nuSVR, phosphorylation propensity from Phospholingo, and net charge per residue.

PLM-derived features included both protein-level and residue-level embeddings. We evaluated embeddings from ESM2 and PTM-mamba. ESM2 embeddings were used to capture latent sequence information at both residue and protein scales, whereas PTM-mamba embeddings were included to represent phosphorylation-aware sequence context but excluded in the final model. In preliminary modeling, four feature configurations were compared: motif-only features, biological features plus motif features, PLM features alone, and a combined representation of PLM and biological features.

### Feature selection and model development

Because the combined biological and PLM feature space was high-dimensional relative to the available sample size, feature selection was performed before final model fitting. We first applied the minimum Redundancy Maximum Relevance (mRMR) method to select features with high relevance to the target variable and low redundancy among themselves. For models using only biological features, 30 selected features gave the best performance. For models using both biological and PLM features, the best first-stage performance was obtained with 25 selected features, and many one-hot encoded sequence features were replaced by embedding-derived dimensions, suggesting that PLM embeddings captured complementary information.

To further refine the feature set, recursive feature elimination with cross-validation (RFECV) was performed. PTM-mamba-derived features were subsequently removed because only two dimensions were retained, and their inclusion would complicate the final prediction pipeline with minimal gain. The final model used 44 features, including 19 ESM residue-level embeddings, 1 ESM protein-level embedding, 19 one-hot encoded amino acid features, and 5 biologically motivated features: condensation propensity, IDR propensity, IDR compactness, phosphorylation propensity, and net charge per residue.

### Modeling method and hyperparameter optimization

The final predictor was implemented using XGBoost through the sklearn interface. XGBoost was selected because previous studies^22^ have shown that gradient-boosted tree models perform well in PTM-site prediction tasks. Hyperparameter optimization was carried out by grid search over key parameters, including n_estimators, learning_rate, max_depth, gamma, reg_alpha, and reg_lambda. The optimal hyperparameter configuration was selected based on cross-validation performance on the training set. The optimized XGBoost model was then used for downstream feature interpretation and independent benchmarking.

### Performance evaluation

To evaluate model performance, five-fold cross-validation was performed on the training set by dividing the data into five non-overlapping subsets. A customized split function was used to ensure that original and augmented data derived from the same protein were assigned to the same subsets. In each round, four subsets were used for training, and one subset was used for validation. Model performance was assessed using precision, recall, F1-score, accuracy, and Matthews correlation coefficient (MCC).

To assess the effect of data augmentation, two baseline models were trained using only one-hot encoded local sequence features from the -7 to +7 phosphosite window: one on the original non-augmented dataset and one on the combined original and augmented dataset. These models were evaluated on both the validation set and the held-out test set.

For comparison with existing tools^1^, we evaluated the final integrated predictor on the independent dataset and compared its performance with a motif-only baseline model trained using the same local-window strategy as previous predictors.

### Model interpretation

To interpret the contribution of individual features to model predictions, we analyzed both the built-in XGBoost feature importance scores and SHAP values. SHAP analysis was used to quantify the effect of each feature on prediction outcomes and to assess the relative contribution of canonical motif features, biological context features, and PLM-derived embeddings in the final model.

## Data availability

The authors provide training data set of 14-3-3 binding proteins and sites as supplement. The human proteome-wide serine/threonine site-level 14-3-3 binding prediction score dataset generated in this study has been deposited in Zenodo under DOI: 10.5281/zenodo.20344516.

## Code availability

All analyses were performed using reproducible notebooks. The complete analysis workflow, including code used to generate the manuscript figures and additional diagnostic, exploratory, and intermediate analysis plots not included in the main or supplementary figures, will be available at https://github.com/liuliu-umich/CAMP-14-3-3. The authors provide the source code and trained models, which will be available at https://github.com/liuliu-umich/CAMP-14-3-3.

## Acknowledgments

This work was supported by American Heart Association (No. 24CDA1272976), University of Michigan Research Scout (No. OORRS033123) and Michigan Biology of Cardiovascular Aging (M-BoCA) Program to LL.

## Author contributions

LL conceived and led the project, designed the methodology, performed the primary experiments, interpreted the results, and wrote the manuscript. ZW provided resources and reviewed the manuscript. XH co-conceived the project, interpreted the results, and edited the manuscript. All authors agreed with the final manuscript.

---

**Supplementary Figure 1.**
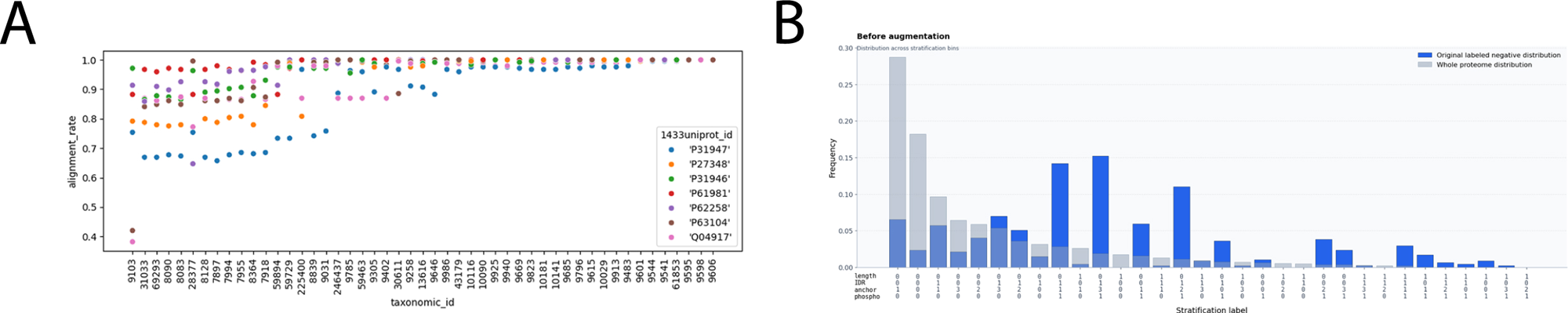
(A) Evolutionary conservation analysis across vertebrate species reveals substantial sequence conservation in 14-3-3 proteins, with alignment rates > 0.6 in 48 species, suggesting functional preservation of binding interactions. (B) Comparison of feature distributions between literature-curated negative sites and the human proteome, showing bias toward shorter, more disordered, and more highly phosphorylatable proteins in the curated negative set before sampling.

**Supplementary Table 1: Data used for this study.**

